# Allatostatin C/AstC-R2 is a Novel Pathway to Modulate Circadian Activity Pattern in *Drosophila*

**DOI:** 10.1101/361048

**Authors:** Madelen M. Díaz, Matthias Schlichting, Katharine C. Abruzzi, Michael Rosbash

## Abstract

Six neuropeptides are expressed within the *Drosophila* brain circadian network. Our previous mRNA profiling suggested that AllatostatinC is a seventh neuropeptide and specifically expressed in dorsal clock neurons (DN1s). Our results here show that AstC is indeed expressed in DN1s, where it oscillates. AstC is also expressed in two less well-characterized circadian neuronal clusters, the DN3s and lateral posterior neurons (LPNs). Behavioral experiments indicate that clock neuron-derived AstC is required to mediate evening locomotor activity under short (winter-like) photoperiods. The AstC-Receptor 2 (AstC-R2) is expressed in LNds, the clock neurons that drive evening locomotor activity, and AstC-R2 is required in these neurons to modulate the same short photoperiod evening phenotype. *Ex vivo* calcium imaging indicates that AstC directly inhibits a single LNd neuron. The results suggest that a novel AstC/AstC-R2 signaling pathway, from dorsal circadian neurons to an LNd, regulates the behavioral response to changing photoperiod in *Drosophila*.

---

Organisms ranging from cyanobacteria to mammals exhibit circadian rhythms or behavioral and physiological processes that occur in a near 24-hour cycle. The molecular clocks that drive these rhythms are sensitive to environmental cues, which provide time-of-day information. Indeed, one important feature of circadian clocks is their plasticity or their ability to re-entrain to different environmental conditions. Clocks can detect fluctuations in light and temperature and adjust features accordingly [1]. This allows adaptation to changing environments, such as seasonal photoperiods and temperatures.

The fruit fly *Drosophila melanogaster* is no exception. Under standard 12:12 light/dark (LD) cycles, it manifests a bimodal locomotor activity pattern with prominent and characteristic morning (M) and evening (E) peaks near the lights-on and lights-off transitions, respectively. Importantly, these peaks show anticipatory behaviors that precede these transitions. Flies also adjust these two locomotor activity events to photoperiod length; this presumably reflects an adaptation to different seasonal conditions. For instance, the M- and E-peaks are further apart under long, summer-like days and closer together under short, winter-like days [2].

These behavioral rhythms are driven by ~150 clock neurons within the fly brain. They are subdivided into distinct neuronal clusters: the small and large lateral neurons (s-LNvs and l-LNvs, respectively), the dorsal lateral neurons (LNds), the lateral posterior neurons (LPNs) and the dorsal neurons (DNs). The DNs are the largest group and are further subdivided into two anterior DN1s (DN1as), sixteen posterior DN1s (DN1ps), two DN2s, and approximately 30-40 DN3s. The s-LNvs are also known as morning cells (M-cells) as they are primarily responsible for the timing of the M-peak, whereas the LNds (along with the 5^th^ s-LNv) are also known as the evening cells (E-cells) as they drive the E-peak [3, 4].

The different clock neuron subgroups must communicate within the circadian network, for example to maintain synchrony of their molecular clocks, especially in the face of varying environmental conditions. Although neuronal communication can occur via gap junctions and neurotransmitters as well as neuropeptides, we have focused here on neuropeptides. To date, only six different neuropeptides have been identified within the circadian neurons. The neuropeptide pigment-dispersing factor (PDF) is expressed in the LNvs, acts as the synchronizing signal to most of the other circadian neurons, and is necessary to maintain rhythmicity in constant darkness [5] [6] [7]. The l-LNvs also express neuropeptide F (NPF) [8], and the s-LNvs co-express short NPF (sNPF) [9]. The activity-promoting E-cells express several different neuropeptides: the 5^th^ s-LNv and five of the six LNds express a combination of NPF, sNPF, and ion transport peptide (ITP) [8] [9] [10]. Among the DN1s, the two DN1as are anatomically and functionally distinct from the DN1ps, partly due to expression of the neuropeptide IPNamide in the DN1as [11]. The neuropeptide diuretic hormone 31 (Dh31) is synthesized in five of the DN1ps to influence sleep [12] and possibly temperature preference [13]. In contrast, there are no neuropeptides associated with any of the remaining circadian clusters and neurons: the DN2s, DN3s, the LPNs, and most of the DN1ps.

To identify additional circadian neuropeptides, we exploited RNA-seq data generated from purified LNvs, LNds, and DN1ps to identify known neuropeptide transcripts that are strongly expressed in circadian neurons [14]. We prioritized the DN1p cluster for the possibility of identifying additional neuropeptides, especially within those neurons with uncharacterized signaling mechanisms. Allatostatin C (AstC) transcripts were highly expressed in the DN1ps and found at much lower levels in the LNvs and the LNds [14]. Intriguingly, transcripts encoding the AstC receptor 2 (AstC-R2) were also detected in the LNds [14]. Thus, we predicted that AstC/AstC-R2 would be a novel intra-clock signaling pathway.

Indeed, AstC is expressed in the DN1s as predicted; it is also expressed in the DN3s and the LPNs. Knockdown experiments indicate that AstC-expressing clock neurons contribute to the advance in evening phase that occurs under a short photoperiod (6:18 LD). Importantly, knock-down of AstC-R2 within the LNds also exhibits the same phenotype as the AstC knockdown, indicating that dorsal neurons communicate with the LNds. *Ex vivo* calcium imaging indicates that AstC directly inhibits a single LNd neuron. These experiments therefore identify a novel AstC/AstC-R2 pathway that regulates evening phase under specific environmental conditions.

## Results

### Identifying AstC as a novel circadian neuropeptide

To characterize gene expression within subpopulations of the *Drosophila* circadian neural network, our lab previously conducted mRNA profiling on the LNvs, LNds, and a subset of the DN1ps (Figure 1*a*), and we focused on neuropeptide transcripts [14]. The sequencing data indicated high levels of the *Allatostatin-C* (*AstC*) transcript in DN1ps, whereas there are lower levels in the LNds (three times lower; *p* < 0.0001) and no detectable *AstC* transcript in the LNvs (Figure 1*b*).

**Figure 1.**
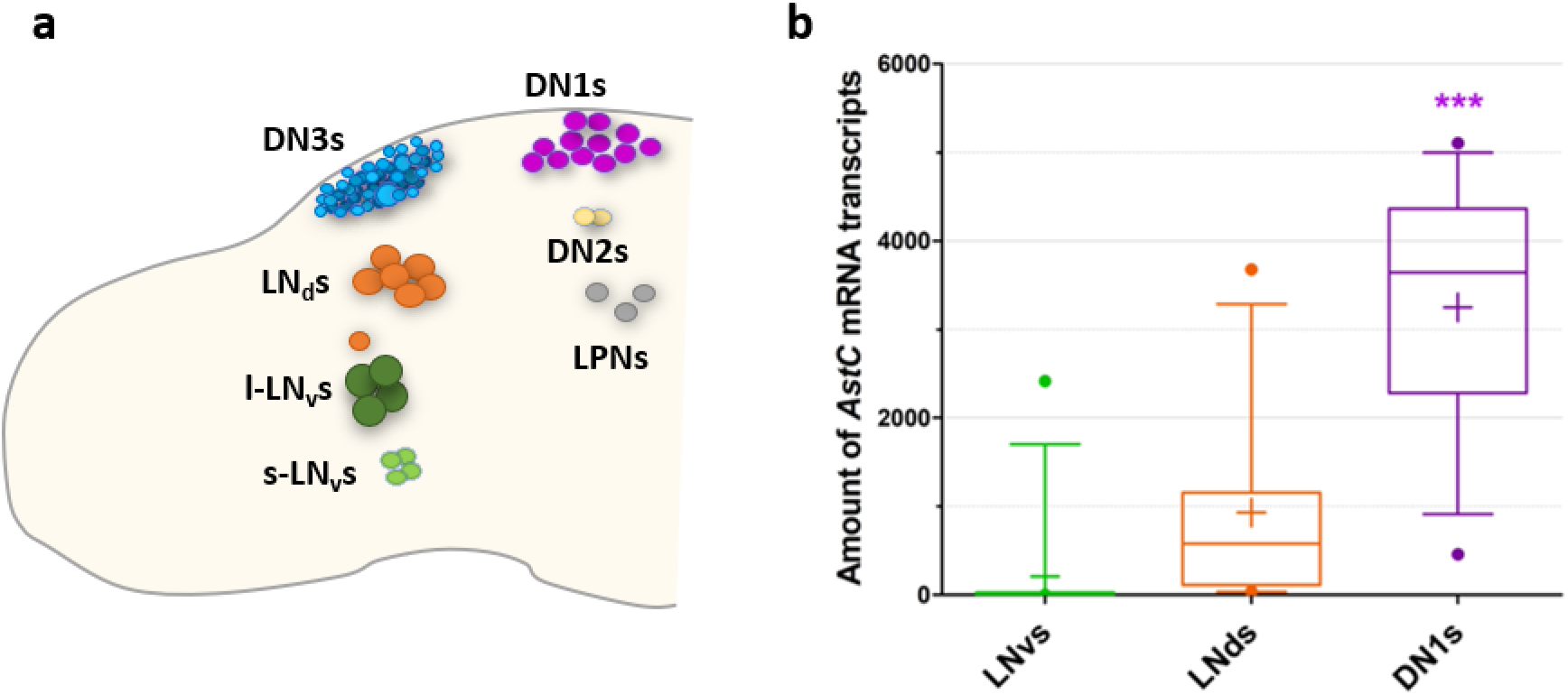
mRNA sequencing data suggests that *Allatostatin C* (*AstC*) mRNA is expressed in the circadian neurons of *Drosophila. **a.*** One hemisphere of the clock neurons in an adult *Drosophila* brain are depicted schematically. The core clock consists of about 150 lateral and dorsal neurons (LNs and DNs, respectively). The ventral LNs are subdivided in the small (s-LNvs) and large neurons (l-LNvs), shown in light and dark green, respectively. The dorsal LNs (LNds) consists of six neurons and the 5^th^ s-LNv, shown in orange. The DNs are subclassified into the approximate 16 DN1s (shown in purple), two DN2s (shown in yellow), and approximately 30-40 DN3s (shown in blue). The three lateral posterior neurons (LPNs) form the last cluster (shown in gray). ***b***. Amount of *AstC* mRNA transcripts in the three neuronal clusters profiled with deep sequencing: the LNvs, LNds (including the 5^th^ s-LNv), and the posterior DN1s. The DN1s have significantly more *AstC* transcripts compared to the LNvs and LNds. Values were averaged from 12 sequencing data sets across different timepoints and biological replicates. Boxplot whiskers show 10^th^-90^th^ percentile. “+” denotes the mean. *** *p* < 0.0001, one-way ANOVA

AstC is known to bind to two G-protein coupled receptors: AstC-R1 (star1) and AstC-R2 (AlCR2) [15]. *AstC-R1* expression was not detectable in the three circadian neuronal groups, but *AstC-R2* transcripts was present in the LNds [14]. Because the transcripts for the peptide and one of its receptors were expressed within circadian neurons, we predicted that AstC plays a role in the clock network and pursued these initial findings.

### AstC is expressed in the DN1ps, DN3s, and LPNs

To confirm that *AstC* transcripts are indeed present in the clock neurons, we took advantage of a recently published fluorescent *in situ* hybridization (FISH) protocol to visualize *AstC* mRNA in whole-mount adult *Drosophila* brains (Figure 2*a*; see materials and methods). The transcripts were detectable in approximately six neurons in the dorsal protocerebrum, a distribution that is quite similar to previously schematized diagrams of *AstC* distribution in adult fly brains [16]. To confirm that these are indeed dorsal circadian neurons, we co-stained with antibody against the core clock transcription factor TIMELESS (TIM). The brains were stained at ZT24, a time when TIM protein is abundant (Figure 2*b*) [17] [18]. Of the five or six AstC-positive neurons in the dorsal brain, only four co-stain with TIM protein (Figure 2*c*), suggesting that this cluster contains four DN1ps, as well as one or two immediately adjacent non-circadian neurons. Surprisingly, *AstC* is also expressed in two additional circadian clusters, the DN3s and the LPNs (Figure 2*c*); these two groups were not profiled [14]. We did not observe *AstC* transcripts in the location of the LNds (data not shown), indicating that the low AstC transcript levels in the LNds indicated by the sequencing is not detectable by FISH or perhaps reflects non-specific background from the RNA sequencing (Figure 1*b*; see Methods).

**Figure 2.**
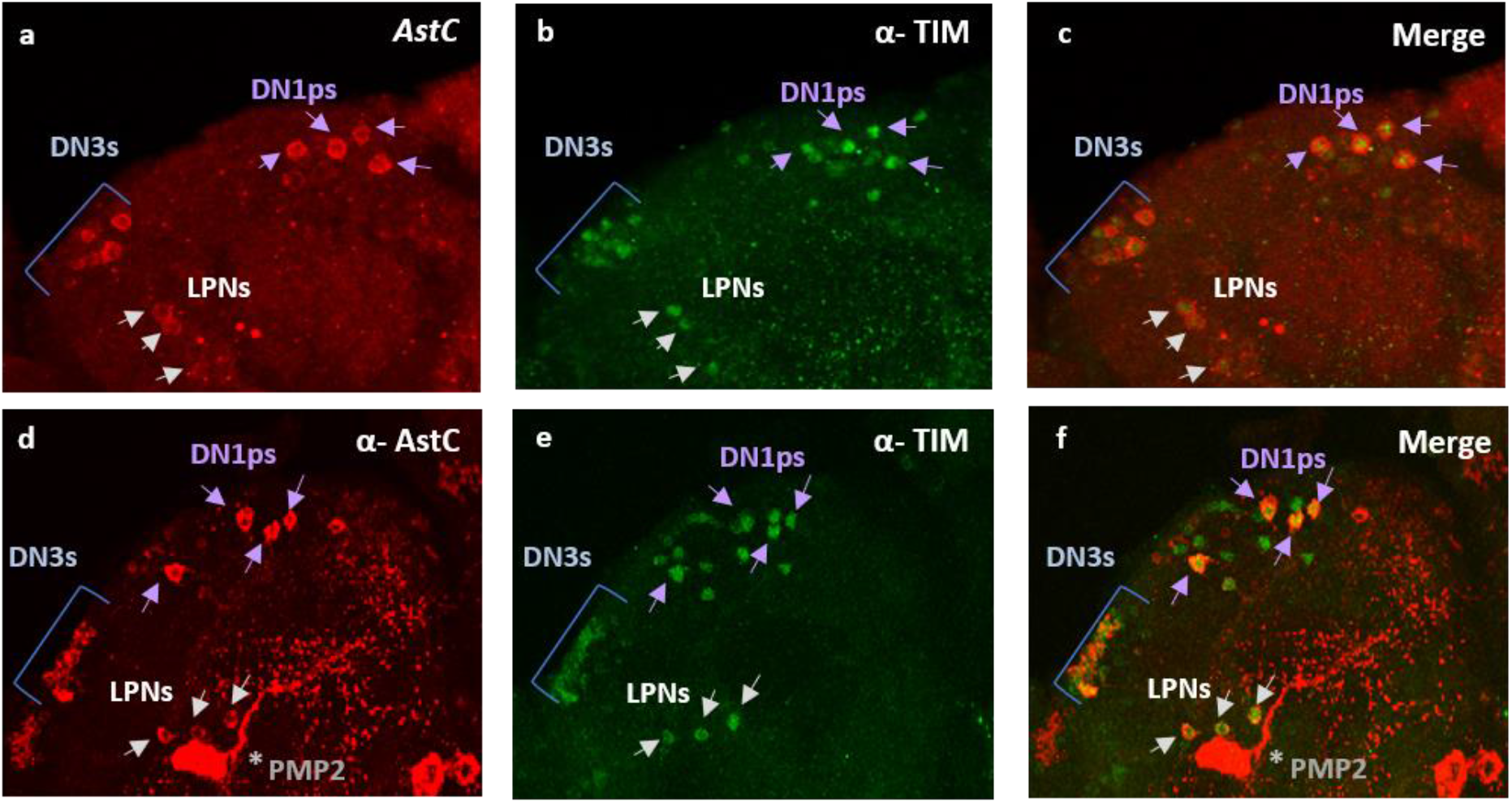
Visualizing AstC in the clock neurons of the adult *Drosophila* brain. ***a***. Fluorescent *in situ* hybridization (FISH) for *AstC* mRNA transcripts at ZT24. ***b***. TIM antibody staining showing the dorsal clock neurons. ***c***. FISH coupled with immunostaining shows the co-localization of *AstC* mRNA transcripts (red) and TIM antibody (green), revealing *AstC* transcripts in four DN1ps (purple arrows), three LPNs (white arrows), and the DN3s (blue bracket). ***d***. Immunostaining of the AstC neuropeptide at ZT20. The posterior medial protocerebral 2 (PMP2, asterisk) are previously known to contain AstC. ***e***. TIM antibody staining showing the dorsal clock neurons. ***f***. Co-localization of AstC neuropeptide (red) is indeed expressed in a subset of the dorsal clock neurons (anti-TIM, green): four DN1ps (purple arrows), three LPNs (white arrows), and the DN3s (blue bracket). Images are from maximum projections. Only the posterior, dorsal region of one hemisphere are shown here. See also Figure S1.

To test whether the *AstC* mRNA is translated into functional peptide within the circadian neurons, we performed co-immunostaining on adult brains during the late night (ZT20) using antibodies against AstC (Figure 2*d*) and TIM (Figure 2*e*). The expression pattern of AstC is very similar to that observed by FISH. The co-localization of AstC and TIM confirms that the AstC peptide is expressed in four DN1ps, approximately 20 of the ~30-40 DN3s neurons, and the three LPNs (Figure 2*f*). AstC is also present within elaborate arborizations in the dorsal regions. Notably, there is a pair of massive non-circadian bi-lateral neurons near the LPNs, which are AstC-positive with likely extensive processes in nearly the entire brain; these cells were previously annotated as posterior medial protocerebral 2 neurons [16, 19] (PMP2; Figure 2*d* and 2*f*, asterisk).

AstC is the first identified neuropeptide in the enigmatic DN3 and LPN circadian clusters. Because previous work from our lab identified the neurotransmitter glutamate within the subset of the DN1ps known to express Dh31 [20], we asked whether AstC is also expressed within this subset of glutamatergic DN1ps. To this end, we expressed GFP with a specific DN1p driver (*R51H05-*GAL4) known to include Dh31 [12] and glutamate [20]. The co-immunostaining with antibodies against AstC and GFP indicated that the four AstC-positive DN1ps are indeed included within this subset that also produces Dh31 and glutamate (Figure S1).

We next asked whether AstC protein levels changes (cycles) in these dorsal clock neurons as a function of time-of-day. We expressed GFP in a subset of the DN1ps and conducted immunohistochemistry co-labeling with anti-GFP and anti-AstC antibodies at six timepoints throughout the 12:12 light:dark (LD) day: ZT0, 4, 8, 12, 16, and 20 (Figure 3).

**Figure 3.**
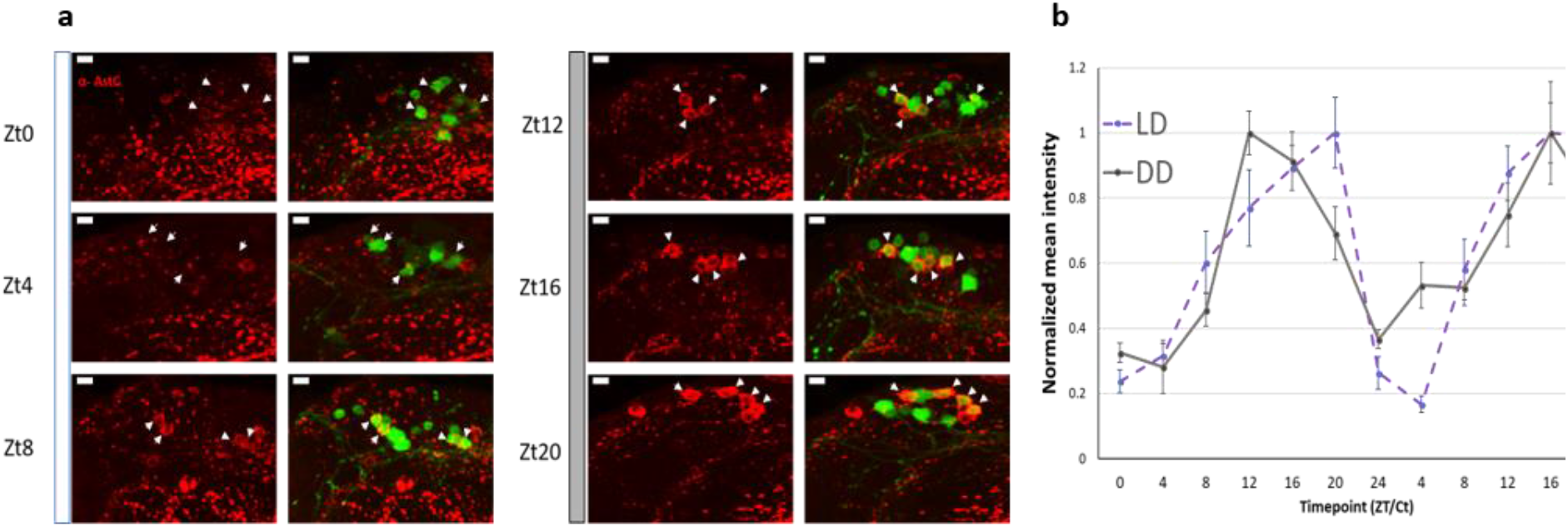
AstC cycles in the DN1ps. Young males flies expressing GFP in a subset of the DN1ps (*clk4.1*-GAL4) were entrained to six timepoints throughout the day under a 12:12 light:dark (LD) cycle. ***a***. Representative images of the DN1ps immunolabeled with anti-AstC (red) and GFP (green). The arrowheads indicate which four AstC-expressing neurons co-localize as the DN1ps. ***b***. Normalized quantification of AstC cycling in the DN1ps under 12:12 LD cycle (dashed, open circles) and the second day of constant darkness (DD, solid, closed circles). AstC is more abundant during the dark phase in comparison to the light phase. Two biological replicates are double-plotted. Error bars are SEM. Images are from maximum projections. *n* ≥5 brains per condition. Scale bar= 5μm. See also Figure S2.

Indeed, AstC cycles throughout the day in the DN1ps: AstC is dramatically reduced during the light phase (between ZT0 and 4), and it accumulates throughout the dark phase, reaching a maximum near ZT20 (Figure 3*b*, *dashed*). To address whether this cycling pattern in the DN1ps is light-driven, we conducted the same immunolabeling experiment in constant darkness (DD) and observed the same DN1p cycling pattern in DD (Figure 3*b*, *solid*), indicating that this is light-independent and therefore almost certainly clock-driven.

The DN3s were identified by their anatomical location. Because AstC can be easily visualized with immunohistochemistry at all time points in LD and DD (Figure S2), we suspect that there is little cycling in this large circadian cluster. However, it is impossible to detect AstC cycling in a subset of the cluster without a stable co-stain like GFP in specific DN3 neurons. The LPNs are also difficult to visualize across timepoints but for a different reason: they are often obscured by the enormous PMP2 neuron that can completely eclipse this clock cluster.

### AstC in the clock neurons contributes to proper timing of the evening phase in a short photoperiod

To explore the function of AstC within the clock network, we knocked-down *AstC* mRNA levels in all clock cells via *tim*-GAL4 driver-mediated expression of an RNAi and then examined the locomotor behavior of these flies under standard 12:12 LD conditions (see materials and methods). We hypothesized that AstC produced in the dorsal neurons may bind to the AstC-R2 in the LNds and affect the evening activity peak (E-peak). However, the behavior of the AstC knockdown and control flies was indistinguishable under these standard conditions (Figure 4*a*). The amplitude and timing of the E-peak were also not significantly different among the genotypes (*p* > 0.05; Figure 4*b*). The negative result was probably not due to inefficient knockdown as indicated by immunostaining of the AstC RNAi strain (Figure S3).

**Figure 4.**
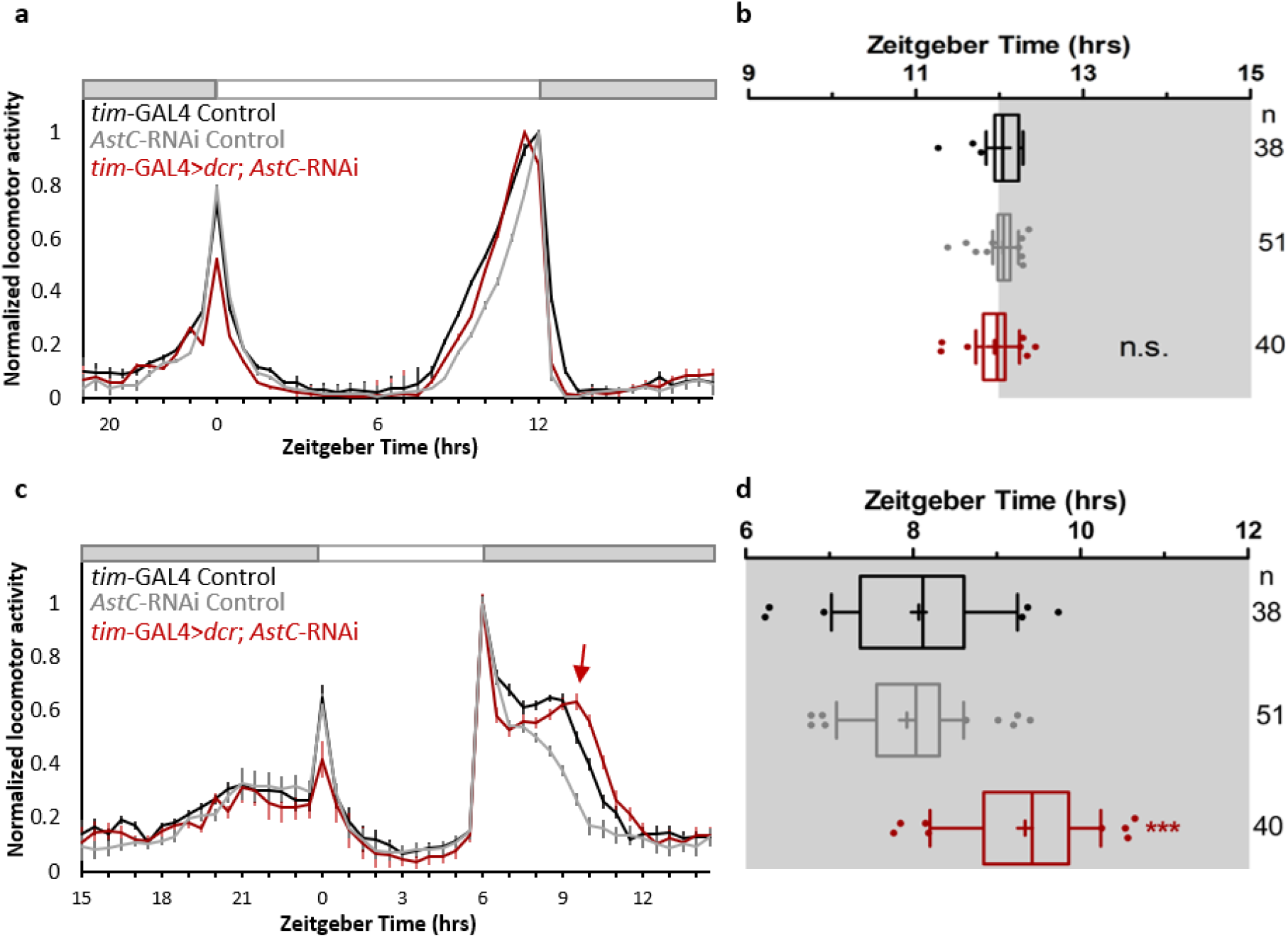
AstC in clock neurons regulates evening phase in short photoperiod days. ***a***. Normalized averaged actograms of the *tim*-GAL4 control (black, n=38), *AstC-*RNAi control (gray, n=51), and the AstC RNAi knock-down in all clock cells mediated by *tim*-GAL4 (red, n=40) under standard 12:12 light:dark (LD) conditions at 27°C. White and dark boxes indicates the respective light and dark phases. The error bars represent SEM. ***b***. Boxplot distribution showing the evening peak phase from individual flies. There are no significant differences (*p* ≥ 0.05). ***c***. Normalized averaged actograms of the *tim*-GAL4 control, *AstC-*RNAi control, and the AstC RNAi knock-down in all clock cells mediated by *tim*-GAL4 under short photoperiods of 6:18 LD at 27°C. The red arrow denotes the delay of the evening peak when AstC is knocked-down compared to the two genetic controls. ***d***. Boxplot distribution showing the evening peak phase from individual flies. The *tim*-GAL4 mediated *AstC-*RNAi knock-down (red) is significantly delayed compared to both the *tim*-GAL4 control (black) and *AstC-*RNAi control (gray; *p* < 0.0001). These data are combined from two independent biological replicates. *** *p* < 0.0001. n.s. no significant difference (*p* ≥ 0.05). “+” indicates the mean and the whiskers denotes the 10^th^/90^th^ percentiles. See also Figures S3, S4, S5, and S6.

Hyperactivity generated by the lights-off event at ZT12 in standard 12:12 LD conditions can override or “mask” more subtle effects on the evening peak [21], [22]. We therefore sought to uncouple the “true” E-peak from this photo-entrainment effect by assaying flies under short photoperiod conditions. To this end, flies were then shifted to 6:18 LD (6 hours light and 18 hours dark) cycles after one week of entrainment under more standard 12:12 LD conditions. Under these 6:18 LD conditions, control flies exhibit an interruption of their day-time siesta by the lights-off transition at ZT6 with the maximum E-peak occurring approximately two hours later at ZT8 (Figure 3*c*, *d*). Flies lacking AstC show a very similar locomotor behavior pattern; however, the E-peak is delayed ~ 1 hour (Figure 3*c; arrow*). Quantification of the maximum evening peak in individual flies showed that there was a significant delay in the timing of the evening peak when AstC was knocked-down in clock neurons (9.34 ± 0.11 hrs) in comparison to the controls (GAL4: 8.07 ± 0.15 hrs; UAS: 7.92 ± 0.09 hrs; *p* < 0.0001; Figure 3*c*, *d*; Figure S4). Because *tim-*GAL4 also drives expression in the eyes, we knocked-down AstC only in the eye with *GMR-*GAL4 driver; there were no significant behavioral differences (*p* > 0.05; Figure S5), strongly indicating that AstC in the brain is responsible for mediating this short period evening locomotor behavior phenotype.

To determine which AstC-positive clock neurons are responsible for this effect, we utilized well-characterized circadian drivers to limit the RNAi expression to more restricted sub-populations. *clk856-*GAL4 targets nearly the entire clock network [23], including the four AstC-positive DN1ps and three LPNS (Figure S6). However, we did not observe significant changes in the evening peak compared to the controls (*p* > 0.05; Figure S5), indicating that AstC from the DN1ps and LPNs is probably not required for mediating the evening phase in short photoperiod conditions. We also did not observe significant differences when AstC was downregulated in the DN1ps with *clk4.1M-*GAL4 (*p* > 0.05; Figure S5), also suggesting that DN1 AstC is not responsible for the phenotype. These negative data suggest that the remaining AstC-expressing DN3s, unlabeled with the *clk856-*GAL4 and *clk4.1M-*GAL4, may be responsible for mediating the short photoperiod evening peak phenotype (Figure S5; Figure S6).

### AstC acts upon AstC-R2 in the LNds

Given the role of the LNds in generating the evening locomotor activity peak [3] [4], we predicted that AstC acts upon the LNds to affect the evening phase short photoperiod phenotype. Of the two receptors, only *AstC-R2* mRNA was detected in the LNds by RNA-sequencing [14]. We therefore knocked-down AstC-R2 in the LNds using *dvPDF-*GAL4, *PDF-*GAL80 mediated RNAi expression (Figure 5). Although the AstC-R2 knock-down was nearly identical to the controls in standard 12:12 LD conditions (Figure 5*a*, *b*), there was a significant delay in the timing of the E-peak in the AstC-R2 knock-down (9.74 ± 0.18 hrs) compared to the genetic controls (GAL4: 8.35 ± 0.13 hrs; UAS: 8.18 ± 0.23 hrs; *p* < 0.0001; Figure 5*c*, *d*) under 6:18 LD conditions. Similar effects were seen using a second and novel split GAL4 driver line (*MB122B-*sGAL4; (Guo, 2017 #9704) [24], which also targets the LNds (Figure S7), i.e., there was a significant delay (*p* < 0.001) in the timing of the E-peak in the knock-down strain (9.05 ± 0.26 hrs) compared to the genetic controls (sGAL4: 7.87 ± 0.10 hrs; UAS: 8.18 ± 0.23 hrs). These phenotypes are essentially identical to those from the AstC knockdowns (Figure 4) and strongly suggest that AstC from dorsal circadian neurons binds to AstC-R2 in the LNds to regulate the timing of the evening peak.

**Figure 5.**
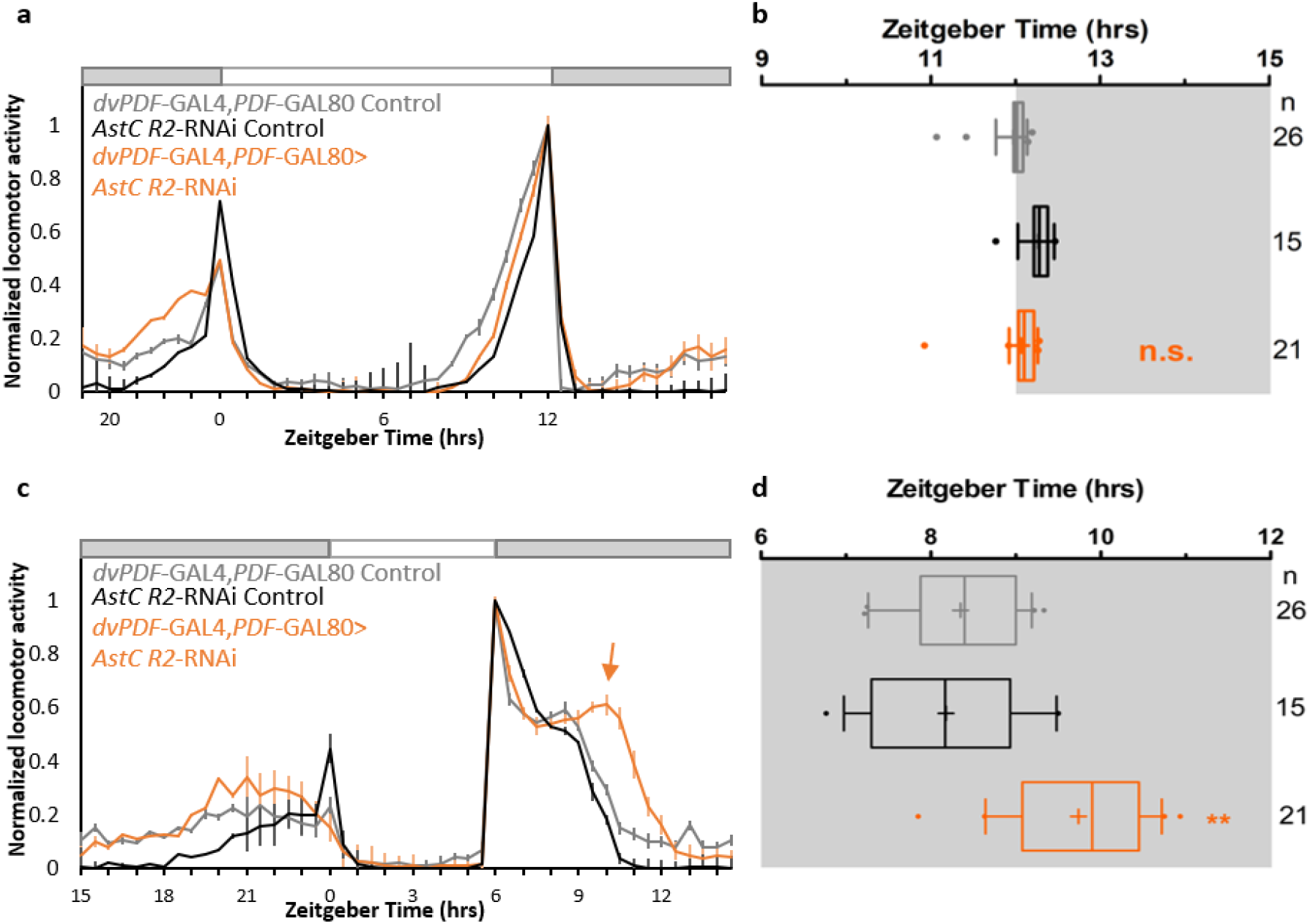
AstC-R2 in LNds is required to regulate evening phase in short photoperiod days. ***a***. Normalized averaged actograms of the *dvPDF*-GAL4, *PDF*-GAL80 control (gray, n=26), *AstC R2-*RNAi control (black, n=15), and the AstC R2-RNAi knock-down in the LNds mediated by *dvPDF*-GAL4, *PDF*-GAL80 (orange, n=21) under standard 12:12 LD condition at 27°C. White and dark boxes indicate the respective light and dark phases. The error bars represent SEM. ***b***. Boxplot distribution showing the evening peak phase from individual flies. There are no significant differences (*p* ≥ 0.05). ***c***. Normalized averaged actograms of the *dvPDF*-GAL4, *PDF-*GAL80 control, *AstC R2-*RNAi control, and the *AstC R2* RNAi knock-down in the LNds mediated by *dvPDF*-GAL4, *PDF*-GAL80 under short photoperiod of 6:18 LD at 27°C. The orange arrow denotes the delay of the evening peak when AstC R2 is knocked-down compared to the two genetic controls. ***d***. Boxplot distribution showing the evening peak phase from individual flies. The *dvPDF*-GAL4, *PDF*-GAL80 mediated *AstC R2-*RNAi knock-down (orange) is significantly delayed compared to both the *dvPDF*-GAL4, *PDF*-GAL80 control (gray) and *AstC R2-*RNAi control (black; *p* < 0.001). ** *p* < 0.001. n.s. no significant difference (*p* ≥ 0.05). “+” indicates the mean and the whiskers denotes the 10^th^/90^th^ percentiles. See also Figure S7.

To address how AstC may modulate the LNds, we conducted functional imaging with the calcium sensor GCaMP6f. Young adult fly brains were collected in the evening between ZT9-12, when LNd calcium is expected to be highest [24, 25]. A baseline period was recorded before exposing the explanted brains to 10μM of synthetic AstC peptide (see methods). We observed significant decreases in fluorescence in a subset of the LNds upon AstC application, indicating that these neurons were inhibited (Figure 6*a*, *b*).

**Figure 6.**
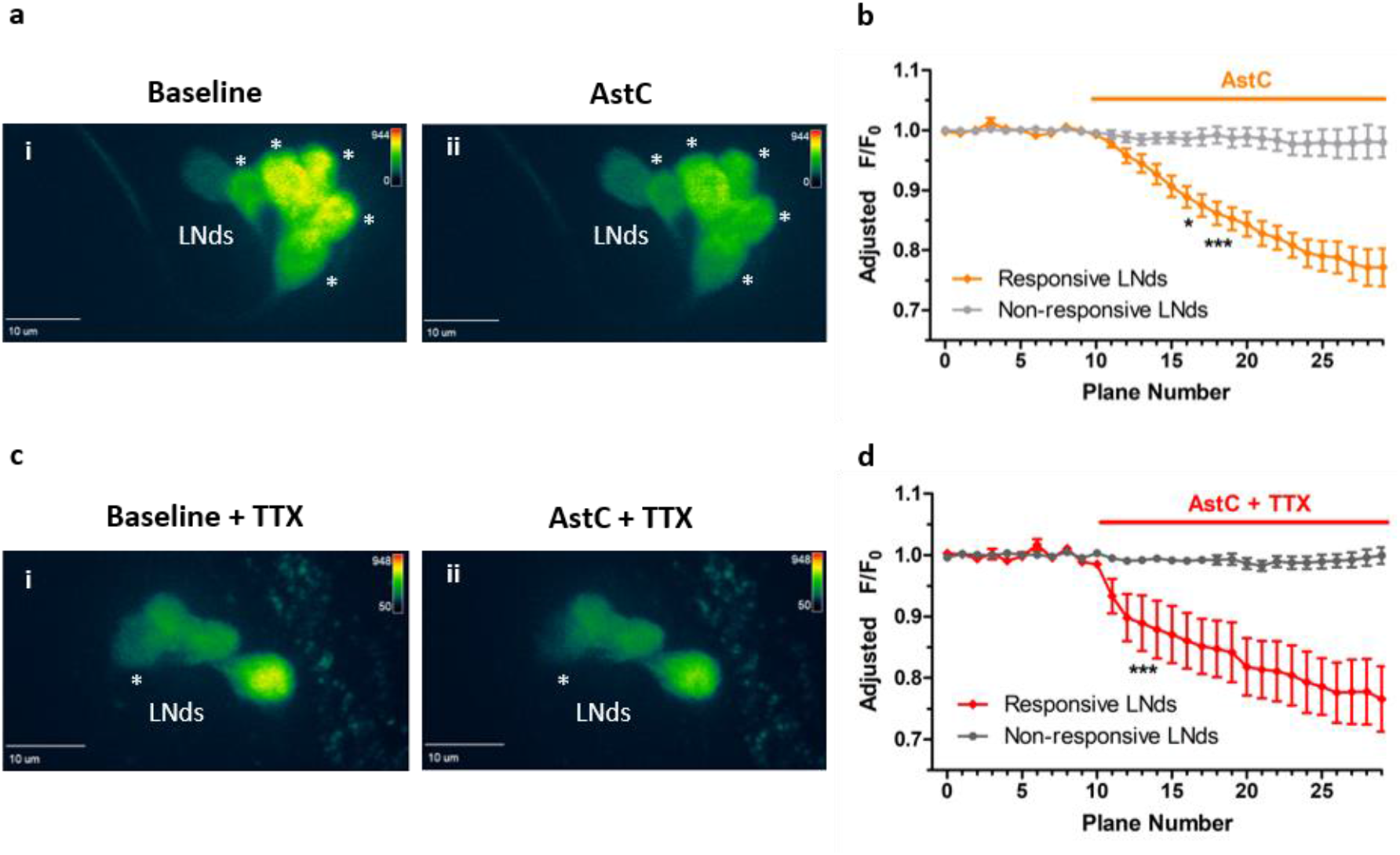
Functional calcium imaging of the LNds responding to AstC (10 μM). Flies expressing the calcium sensor GCaMP6 in most clock neurons (*clk856*-GAL4> UAS-GCaMP6f) were collected in the evening, ZT9-12, and brains were explanted. ***a***. A baseline recording of the LNds was obtained for 10 planes (Ai). When the synthetic AstC peptide (10 μM) was added into the bath, a substantial decrease in fluorescence was observed in multiple LNd neurons (Aii, denoted by asterisks). ***b***. Quantification of the fluorescence relative to the initial baseline recording (F/F_0_) over time. The LNd neurons showed either a significant decrease in calcium (orange, ‘Responsive LNds’) or were unaffected (light gray, ‘Non-responsive LNds’). ***c***. To address whether AstC directly inhibited the LNds, the same experiment was conducted with the sodium channel blocker tetrodotoxin (TTX) added to the bath. Similar to A, a baseline recording was obtained for the LNds (Ci) before exposing the brains to AstC (Cii). The asterisk denotes the single LNd neuron that showed a significant decrease in calcium in response to the AstC treatment. ***d***. the quantification of the direct inhibition of AstC onto a single LNd neuron (red, ‘Responsive LNds’). Most LNd neurons were unaffected. *n* = 6 brains per condition. * *p* < 0.05, *** *p* <0.001, two-way ANOVA.

Because the number of inhibited LNds neurons varied widely from only one to five, we were concerned about indirect network interference. To address the extent to which AstC directly inhibits the LNds, the same functional imaging experiments were conducted in the presence of tetrodoxin (TTX), a voltage-gated sodium channel blocker. Similar decreases of LNd fluorescence signal were observed (Figure 6*c*, *d*), indicating that AstC indeed directly inhibits the LNds. However, there was much less variability in the number of responsive LNds in TTX. Although most of the LNds showed no significant changes compared to baseline recording upon AstC application (Figure 6*d*, gray), a single LNd neuron was consistently responsive with a dramatically decreased calcium signal (Figure 6*c*ii, asterisk; 6*d*, red). These data indicate that AstC directly inhibits a single LNd neuron, indicating striking specificity within a substantially heterogeneous clock neuronal cluster.

## Discussion

To learn more about how the ~150 clock neurons within the adult fly brain communicate, we examined RNA sequencing data from the LNds, LNvs and DN1s for neuropeptides not yet associated with this circuitry. AstC was a promising candidate because mRNAs encoding both the peptide and one of its receptors (AstC-R2) were identified within the three clock neuron clusters; these data suggested a novel intra-clock circuitry signaling pathway. *AstC* transcripts as well as the neuropeptide are indeed well-expressed in the DN1s, and the neuropeptide signal undergoes strong cycling in DD as well as LD conditions. Moreover, AstC is also expressed in two other circadian neuron subgroups, the DN3s and the LPNs, and it is the first neuropeptide identified in these circadian clusters. Behavioral data after RNAi knockdown experiments indicate that the AstC binds to AstC-R2 expressed in E-cells to modulate the timing of evening locomotor activity. *Ex vivo* calcium imaging indicates that AstC directly inhibits a single LNd neuron.

### Functional role of AstC and AstC-R2 in clock neurons

AstC is required in the clock neurons to regulate the evening locomotor activity phase in short photoperiods. The shift in the timing of the evening peak occurs when AstC is reduced in all circadian neurons (*tim*-GAL4 driver), and the DN1 cycling pattern suggests a circadian modulation of secretion. Although this temporal regulation could coincide with the timing of its effect on the phase of the evening locomotor peak, reducing AstC solely in the DN1ps using *clk4.1M-GAL4* was without effect (Figure S5). The LPNs are also probably not a key circadian source of AstC: their AstC levels were dramatically reduced in the *clk856-*GAL4 mediated knock-down, yet no phenotype was observed (Figure S5). All AstC-expressing DN3s are targeted by the *tim*-GAL4 driver, and most of these DN3s are not included in the *clk856-*GAL4 driver (Figure S6). These results suggest that the DN3s may be the source of AstC that is critical for the differences in evening locomotor activity. Unfortunately, this tentative conclusion is based on negative data, and the lack of a DN3-specific GAL4 driver makes it impossible to test this model directly. Therefore, we propose three possible models: 1) the DN3s are the primary source of AstC within the circadian circuit (Figure 7*a*); 2) the DN1s, DN3s, and LPNs, or some combination, are functionally redundant (Figure 7*b*); 3) A small amount of residual AstC within DN1s is sufficient for its behavioral role in the evening activity peak assay. Although neuron-specific deletion of *AstC* would contribute to distinguishing between these three possibilities, we do not have an accurate and efficient CRISPR-based strategy for achieving temporal and spatial specificity.

**Figure 7.**
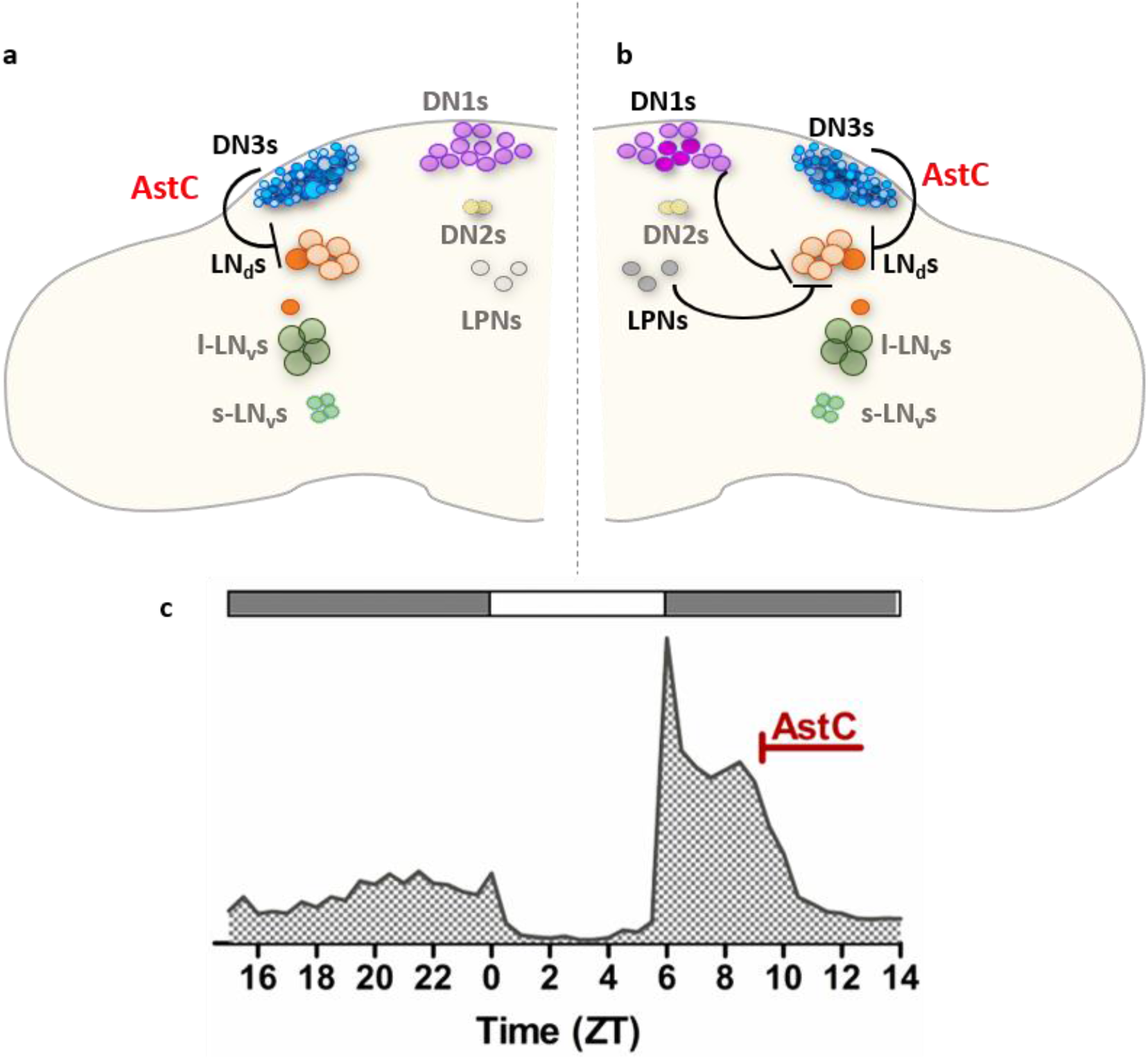
Proposed model for AstC/AstC-R2 signaling in the circadian circuitry of *Drosophila*. ***a***. The DN3s are the integrator for light/dark duration to sense changes in photoperiod conditions. AstC solely in the DN3s inhibits a single LNd neuron via AstC-R2 to regulate the evening phase, as revealed under short photoperiod. ***b***. It is unclear which group of clock neurons serves as the critical integrator of the different photoperiod conditions. Due to lack of a DN3-specific driver, we propose this second model: a combination of the AstC-containing DN1ps, DN3s, and LPNs, together inhibit an LNd neuron to regulate the evening peak, indicating a functional redundancy in neuropeptide signaling. ***c***. The functional role of AstC is to precisely regulate the evening phase, as seen under short photoperiod condition. Because the reduction of AstC leads to a delay in the evening phase, AstC must normally act to regulate the locomotor evening offset.

Once released by dorsal circadian neurons, AstC signals to the LNds via binding to its receptor, AstC-R2. This is because AstC-R2 knock-down in the LNds and knockdown of AstC in the entire circadian circuit give rise to the identical phenotype, a delayed evening peak in short photoperiod conditions (Figure 4, Figure 5, Figure S7). Moreover, the DNs and the LNds both extend projections to the dorsal protocerebrum region (near the pars intercerebralis), where they come in close proximity [26], which would facilitate neuropeptide transmission.

Our functional imaging strongly indicates that AstC binding to LNd-localized AstC-R2 leads to neuronal inhibition. The effect is consistent with several previous electrophysiology experiments [15, 27, 28] and contributes to an emerging theme of LNd inhibition [20, 24, 25, 29]. It is also one of the first pieces of evidence indicating that the dorsal neurons can be a source of inhibition onto the LNds [20]. Interestingly, only a single LNd neuron is directly AstC-sensitive, further attesting to LNd heterogeneity [9] [8] [10] and suggesting that behavioral regulation of the evening phase under short photoperiod arises from this signaling to a single LNd neuron. The response of that neuron may be communicated to the rest of the LNds, for example via gap junctions.

Previous studies provide hints that the DN3s could inhibit the LNds. For example, the timing of the calcium phase in DN3s, a surrogate of neuronal activity, is coincident with a decrease in LNd calcium [24]. The same reciprocal relationship was also observed under short photoperiods. Under these conditions, the DN3s manifest an increase in calcium signaling immediately after the lights-off transition. This occurs immediately after the actual E-peak of locomotor activity and coincides with a calcium decrease in the LNds [25]. Although more experiments are needed to support this hypothesis, it is possible that the DN3s may inhibit this single LNd via AstC release to modulate the proper timing of evening locomotor activity (Figure 7*a*).

There are several caveats to this current model. First, due to the lack of functional RNAi lines available, a single RNAi line was used for both the AstC and AstC-R2 knockdown experiments. Although not ideal, this concern is partially alleviated by the observation that the AstC and AstC-R2 knockdowns show essentially identical phenotypes suggesting that off-target effects are only minimally relevant. Second, we have been unable to rule out the possibility that the phenotypes observed in AstC knockdowns are not due to a requirement for AstC during development. Experiments to address this point are challenging because when the temperature is raised to reduce tubgal80^t^s repression and allow for adult-only knockdown, the heat itself drastically changes the timing of the evening peak. Third, lack of a DN3-specific driver has precluded determining whether DN3-derived AstC is required for this evening activity peak modulation, and whether the DN3s can directly inhibit the LNds.

### A conserved role of allatostatin C/ mammalian somatostatin in photoperiodism

Although the AstC peptide sequence is highly conserved among insect species [30] [31] [32], only the AstC-R2 receptor has a mammalian homolog, the somatostatin/galanin/opioid receptor family [15, 27, 28]. Inhibitory somatostatin (SST) interneurons are present in the mammalian equivalent of a central core clock, the suprachiasmatic nucleus (SCN) [33] [34] [35]. SST interneurons are also known to affect sleep [36] and circadian behaviors [34]. Interestingly, SST is associated with proper adaptation under short-day conditions for both diurnal and nocturnal mammals [37-39], suggesting a highly conserved function with AstC/AstC-R2 for adaptation under different seasonal environments. It will be interesting to see whether the SCN-resident SST interneurons are important for this adaptation, like the AstC-containing clock neurons described in this study.

## Methods

### *Drosophila* stocks

*D*. *melanogaster* strains were reared on standard cornmeal/agar medium supplemented with yeast under 12:12 LD cycles at 25 °C. The following transgenic flies were used for behavior: *Tim-*GAL4 (yw; *Tim-*GAL4/CyO) [40], *Clk4.1M-*GAL4 [41], *Clk856-*GAL4 [23], *GMR-*GAL4 [42], *dvPDF-*GAL4 [43], *PDF-*GAL80 [4], and UAS*-Dicer2*. The LNd *split-*GAL4 *(JRC_MB122B)* was provided by Gerald M. Rubin. UAS*-AstC* RNAi and UAS*-AstC-R2* RNAi lines (nos. 102735KK and 106146KK, respectively) were from the Vienna Drosophila RNAi Center. The GAL4 controls were crossed into the empty background vector of the RNAi (60100KK). Microscopy experiments involved *W^1118^* for wild-type flies, *R51H05*-GAL4 [44], and UAS-*EGFP*. Young, male flies were used for all experiments.

### Quantification of mRNA sequencing data

mRNA profiling of specific clock neurons was previously described in [14, 45]. For the distribution of the differential sequencing of *AstC*, the mRNA transcripts were averaged across the entire *AstC* gene for each of the twelve data sets. The twelve independent data sets originate from two biological replicates of each of the six time-of-day collections. Statistical analysis was performed on the Prism 5 software (GraphPad) using a one-way ANOVA with a Tukey’s multiple-comparisons test.

### Fluorescent *in situ* hybridization (FISH)

FISH was performed as described previously [46] onto wild-type flies (*W^1118^*) at ZT 24 with the following exceptions: custom oligo probes were ordered against the entire *Ast-C* mRNA sequence, including the 5’ and 3’ untranslated regions, and conjugated with Quasar 670 dye (Stellaris Probes, Biosearch Technologies). For the hybridization reaction of the probes onto the brains, the oligo probes were diluted to a final concentration of 0.75 μM. Immediately following the FISH protocol, the brains were blocked with 10% normal goat serum for two hours at room temperature before incubating in primary antibody overnight at 4°C (α-TIM 1:200). The brains were then fluorescently labeled with Alexa Flour 488 conjugated anti-rat at 1:500 for four hours at room temperature. Lastly, the brains were washed before mounting onto slides with Vectashield Mounting Medium (Vector Laboratories). The slides were immediately viewed on a Zeiss 880 series confocal microscope with a 25x oil objective. The z-stack was sequentially imaged in 0.8 μm sections.

### Adult fly brain immunohistochemistry

Wild-type flies (*W^1118^*) were entrained for three days before collecting at their respective timepoints. Fly heads were fixed in PBS with 4% paraformaldehyde and 0.008% Triton X-100 for 60-65 min at room temperature while rotating. Fixed heads were washed in PBS with 0.5% Triton X-100 (PBS-T) and then dissected in PBS-T. The brains were blocked in 10% normal goat serum (NGS; Jackson Immunoresearch) for an hour at room temperature. The brains were later incubated with primary antibodies at 4 °C for three nights. For TIM and AstC co-staining, rat anti-TIM antibody (1:200) and rabbit anti-*Manduca* AstC antibody (1:250, gift from Dr. Jan Veenstra) were used as primary antibodies. After three washes with PBST, the brains were incubated with Alexa Fluor 488-conjugated anti-rat and Alexa Fluor 635-conjugated anti-rabbit (Molecular Probes) at 1:500 dilutions in 10% NGS. After the brains were washed three more times, the brains were mounted in Vectashield Mounting Medium (Vector Laboratories). The slides were immediately viewed on a Leica SP5 confocal microscope with a 20x objective and sequentially imaged in 0.8 μm sections. For experiments showing the validation of the RNAi efficiency, the laser intensity and other settings were set the same across the different genotypes for each experiment.

### Locomotor behavior assay

We used the *Drosophila* Activity Monitoring system (Trikinetics, Waltham, MA, USA) to record the number of beam crosses caused by the fly in one-min intervals. Young, male flies were individually placed in glass tubes containing agar-sucrose media. The temperature was constant at 27°C for increased knock-down efficiency. Flies were allowed one day of habituation before six days of entrainment under 12:12 LD. For the determination of the evening peak timing under short days, the flies were then transferred to 6:18 LD for six additional days. Only the final four days of locomotor activity were used in our analysis. Average activity recorded from short photoperiod day 3–6 was plotted for each fly and the average evening peak time scored as described previously [47]. Statistical analysis was performed on the Prism 5 software (GraphPad) using a one-way ANOVA with a Tukey’s multiple-comparisons test.

### Functional calcium imaging

The calcium sensor GCaMP6f was expressed in most clock neurons using *clk856*-GAL4. Young male flies were entrained to a standard 12:12 LD to be collected at evening time between ZT9-12. Adult male fly brains were dissected in adult hemolymph-like saline (AHL) (108 mM NaCl, 5 mM KCl, 2 mM CaCl_2_, 8.2 mM MgCl_2_, 4 mM NaHCO_3_, 1 mM NaH_2_PO_4_ -H_2_O, 5 mM trehalose, 10 mM sucrose, 5 mM HEPES, pH 7.5; Wang, 2003 #6627). Brains were then pinned to a layer of Sylgard (Dow Corning, Midland, MI) silicone under a small bath of AHL contained within a recording/perfusion chamber (Warner Instruments, Hamden, CT, RC-26G) and bathed with room temperature AHL. Brains expressing GCaMP6f were exposed to the fluorescent light for approximately one to two minutes before imaging to allow for baseline fluorescence stabilization of the LNds. Perfusion flow was established over the brain with a gravity-fed ValveLink perfusion system (Automate Scientific, Berkeley, CA). A spinning disk confocal (Intelligent Imaging Innovations, Inc., Denver, CO) was used to visualize all the LNds possible by recording a z-stack with a step size ranging from 2-2.5 μm. After 10 planes of baseline recording with AHL, 10 μM AstC (GenScript, Piscataway, New Jersey) was delivered by switching the perfusion flow until the end of the recording for a total of 30 planes.

For tetrodoxin (TTX) experiments, brains were dissected with regular AHL before being pre-incubated in AHL containing 1 uM TTX for at least 5 minutes. The brains were then recorded in the same parameters as above, with AHL+TTX (1 uM) as a baseline recording before exposing the brains to AstC (10 uM) + TTX (1 uM).

The Slidebook Reader software was used to subtract the background from the signal intensities of the regions of interest. After the background subtraction, the initial F/F_0_ values were exported. Based on the 10 planes of baseline recording, a bleach trend line was calculated for each sample (MATLAB 2017, MathWorks, Natick, MA), and therefore the F/F_0_ was adjusted to reflect any true fluorescent signal changes. Individual LNd neurons were considered ‘responsive’ if there was at least a 10% change in fluorescence signal. Statistical analysis was performed on the Prism 5 software (GraphPad) using a two-way ANOVA.

## Acknowledgements

Many thanks to members of the Rosbash lab for several thoughtful discussions, and especially Dr. Fang Guo, Meghana Holla, and Patrick Weidner for insightful comments. Much appreciation to members of the Griffith Lab at Brandeis University, especially Johanna “Joey” Adams and Dr. Timothy Wiggin for post-hoc analysis of the functional imaging. Also, Muibat Yussuf for assistance and fly maintenance. Thanks to Dr. Orie Shafer for technical guidance. Thank you to Drs. Xi “Salina” Long, Robert Singer, and Timothee Lionnet for technical assistance with the FISH protocol and to Ed Dougherty for microscopy support. The AstC antibody was a generous gift from Dr. Jan Veenstra. We thank the Bloomington Stock Center and Vienna Drosophila RNAi Center for flies.

## Author contributions

Conceptualization, M.D., M.S., K.A., and M.R.; Methodology, M.D and M.S.; Formal Analysis, M.D. and M.S.; Investigation, Visualization, and Writing-Original draft, M.D.; Writing-Review and Editing, M.S., K.A., and M.R.; Project Administration, M.S., K.A., and M.R.; Funding acquisition, M.R.

## Competing interest

The authors declare no conflicts of interest.

## Supplemental figures

**Figure S1-.**
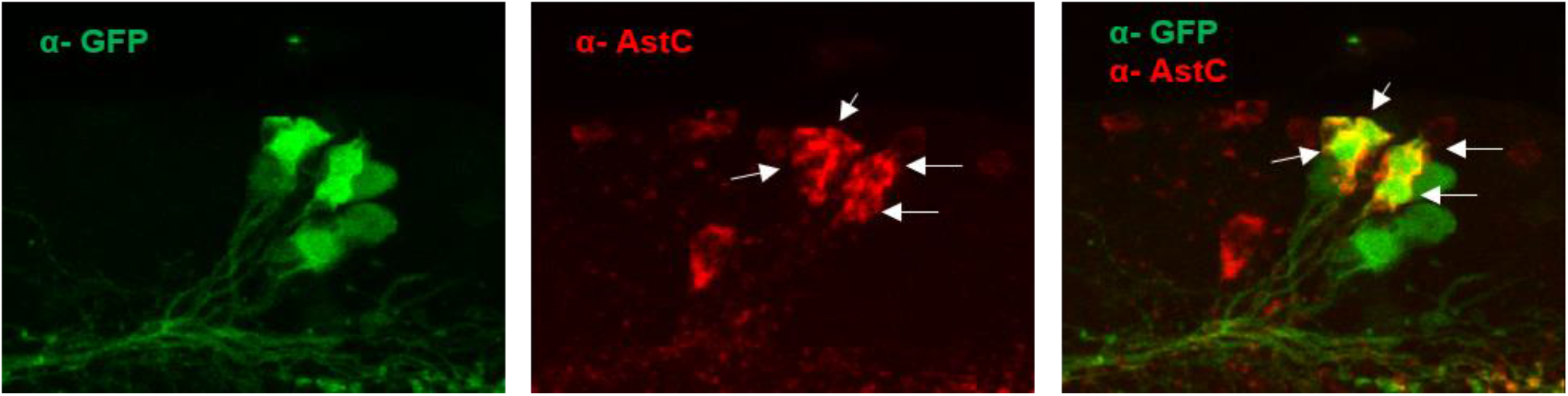
AstC is found in glutamatergic DN1ps. Related to Figure 2. A subset of the DN1ps known to contain glutamate (*R51H05*-GAL4) was labeled with GFP and co-immunostained with AstC antibodies at ZT14. Co-localization of the two signals indicates that AstC is found in four glutamatergic DN1ps. Non-clock neurons also expressing AstC are found nearby.

**Figure S2-.**
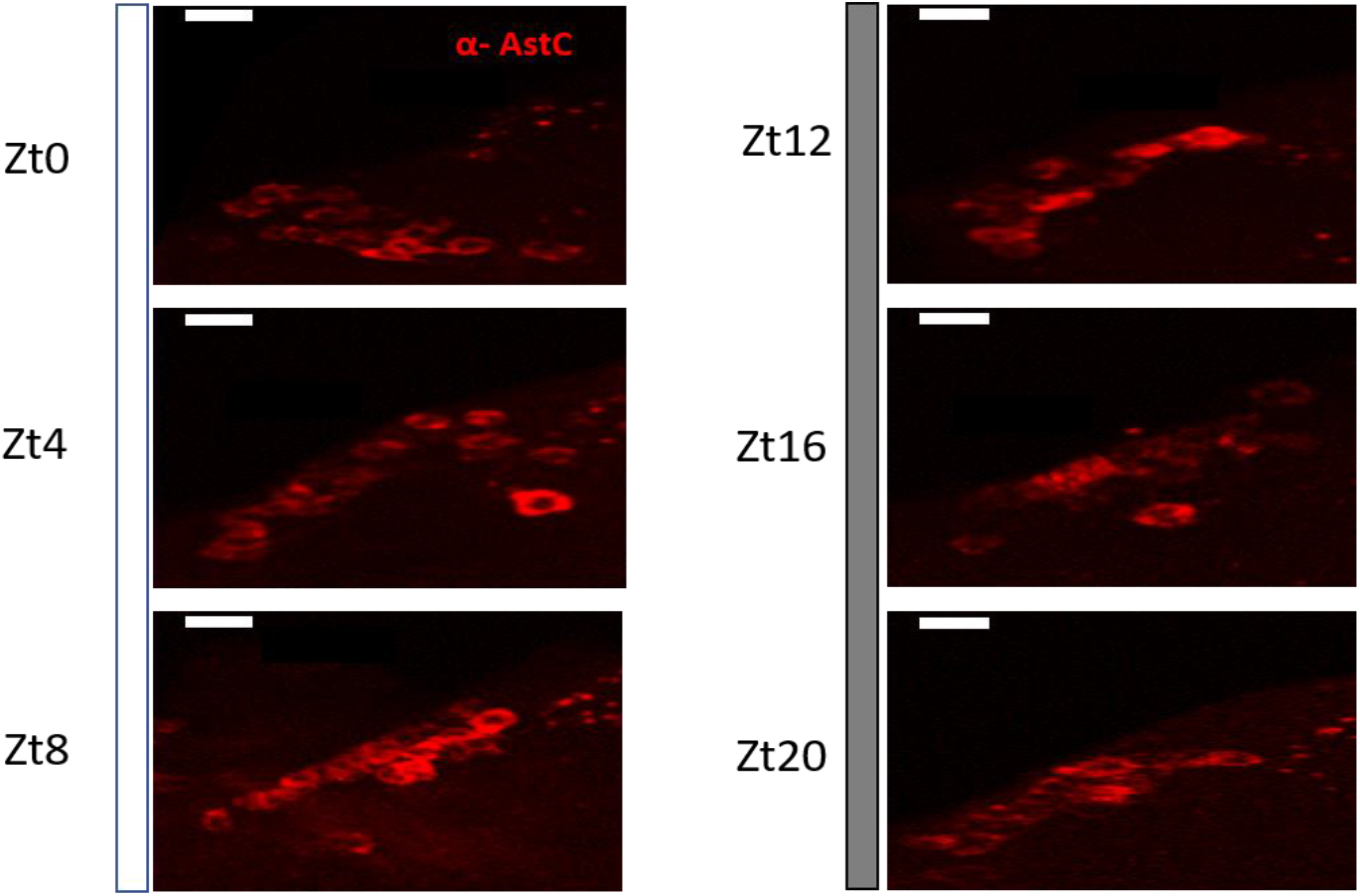
AstC in the DN3s can always be visualized across timepoints. Related to Figure 3. Flies were collected across six timepoints in a standard 12:12 LD cycle. Representative images of the DN3s show that AstC is always visible in the cluster. Scale bar= 5μm.

**Figure S3-.**
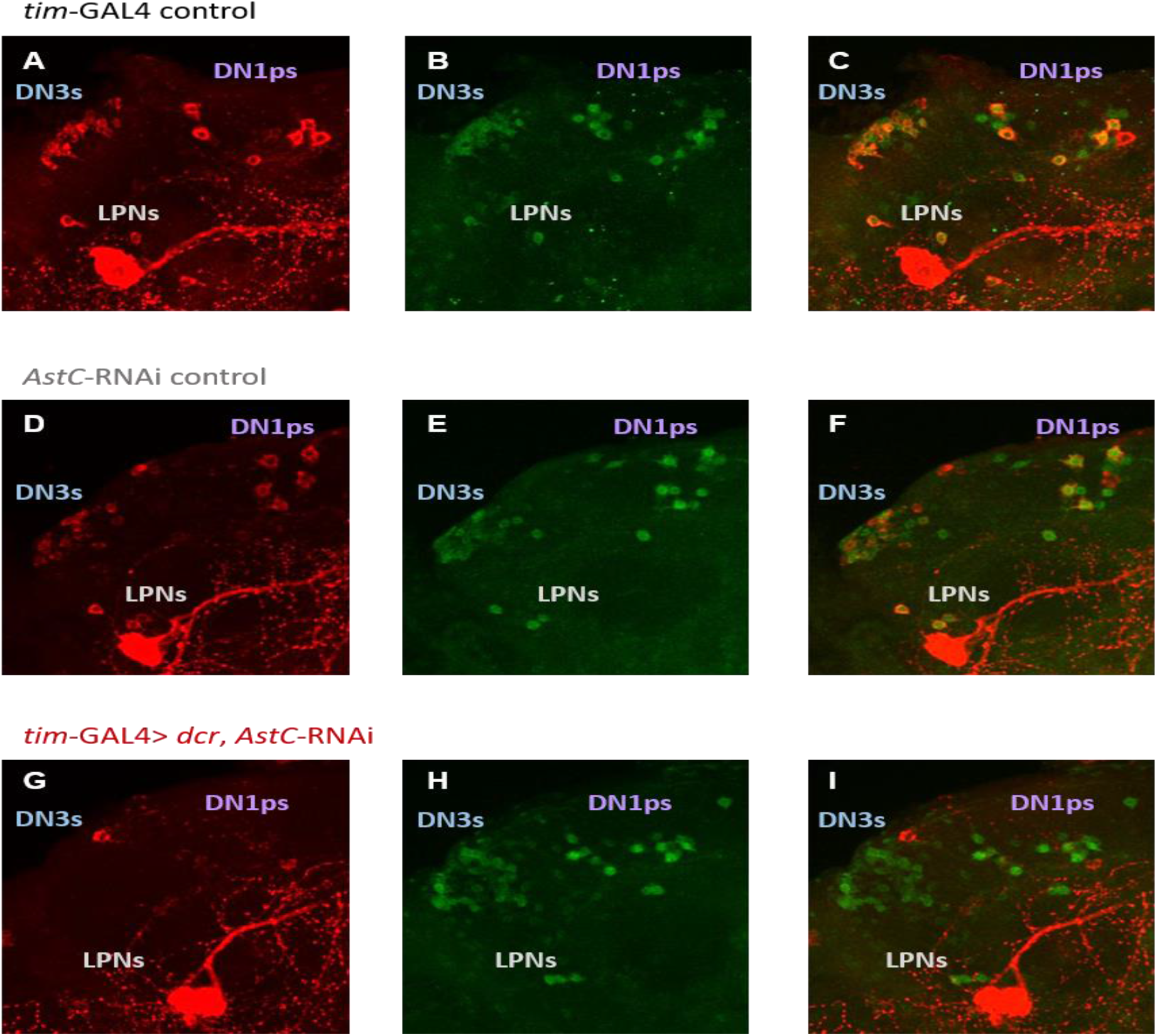
Immunostaining validation of AstC RNAi knock-down in all clock cells. Related to Figure 4. *AstC-*RNAi was expressed with *tim*-GAL4>*dcr* to efficiently knock-down AstC peptide in all clock cells. The GAL4 control (A-C), UAS control (D-F), and the experimental knock-down genotype (G-I) were stained with anti-AstC (red) and anti-TIM (green) at ZT20. ***A***-***F***, For both parental controls, AstC is expressed in four DN1ps, approximately half the DN3s, and the three LPNs. ***G***-***I***, AstC expression is abolished in all clock neurons in the experimental knock-down group. At least eight brains of each genotype were stained together under the same conditions. Confocal settings were maintained the same for all genotypes.

**Figure S4-.**
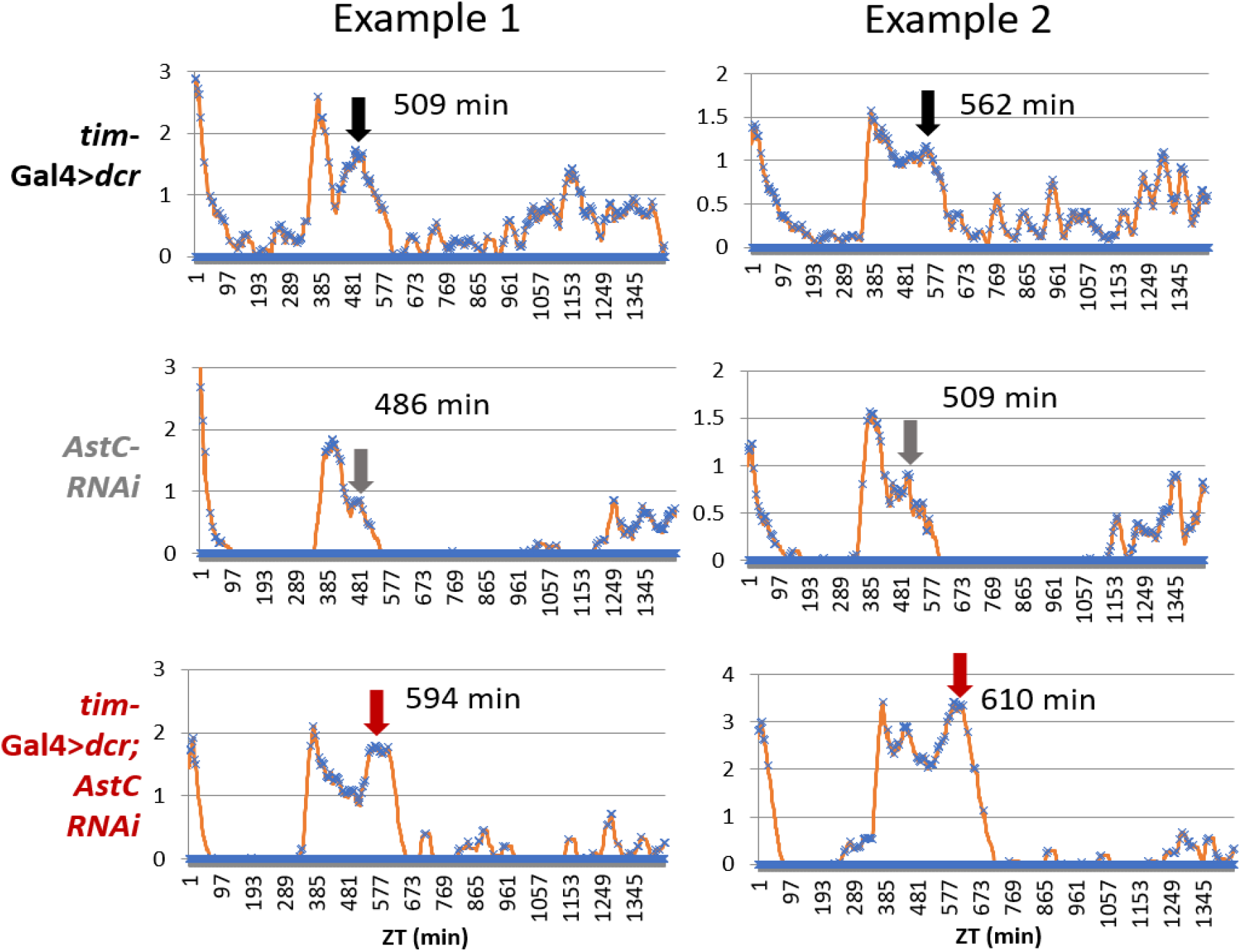
Representative actograms in 6:18 LD. Related to Figure 4. Averaged actograms from two individual flies are shown for the *tim*-GAL4 control (top, black), the *AstC*-RNAi control (middle, gray), and the experimental group with AstC knocked-down in all clock neurons (bottom, red). The arrows indicate the timing of the maximum E-peak considered for analysis. Actograms averaged from Days 3-6 in 6:18 LD. Y-axis measures total activity counts per minute.

**Figure S5-.**
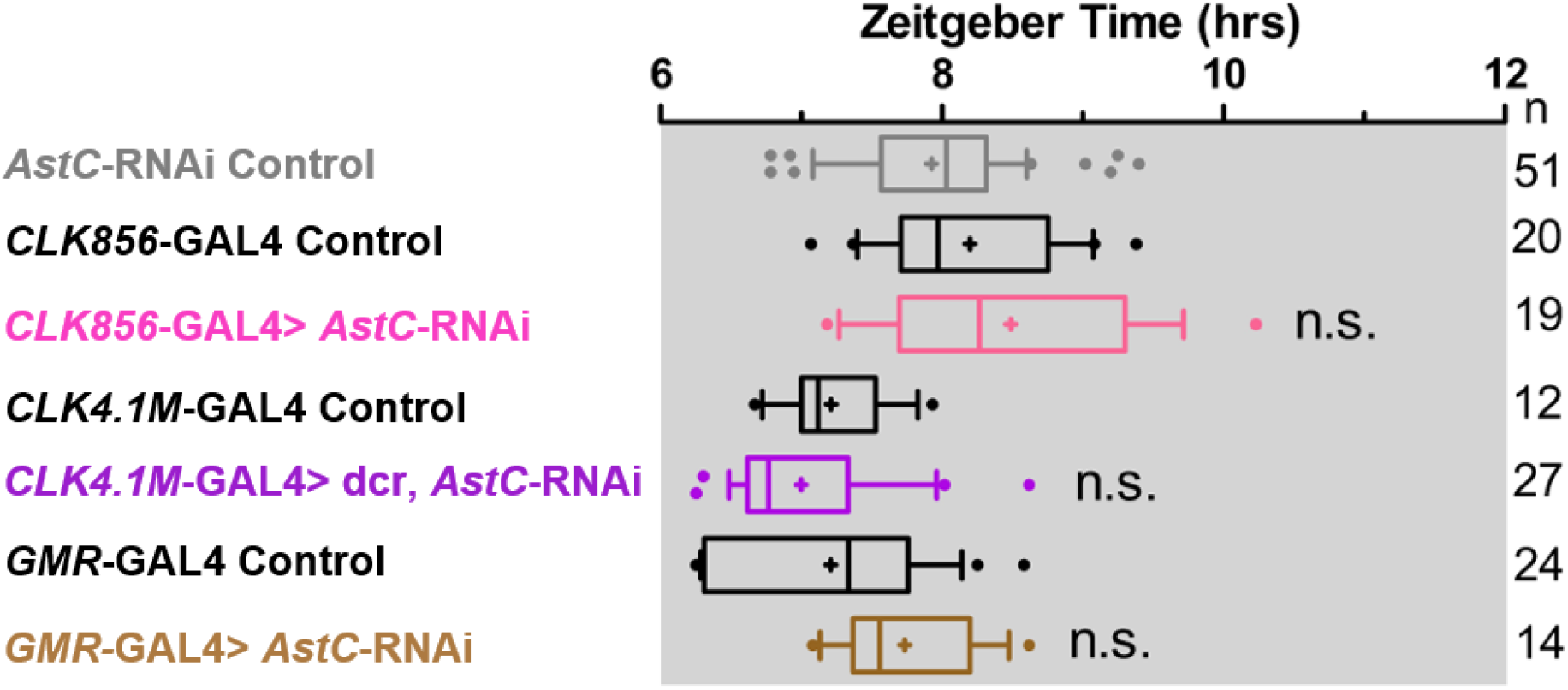
AstC knock-down in subsets of clock neurons has no significant effects in E-peak under short photoperiod. Related to Figure 4. Boxplot distribution showing the evening peak phase measured from individual flies. The *clk856*-GAL4 mediated *AstC-*RNAi knock-down (pink) is not significantly different to both parental controls. AstC knock-down mediated by *CLK4.1M*-GAL4 (purple) and *GMR*-GAL4 (brown) also showed no significant differences compared to their respective parental controls. n.s. no significant difference (*p* ≥ 0.05). “+” indicates the mean and the whiskers denotes the 10^th^/90^th^ percentiles.

**Figure S6-.**
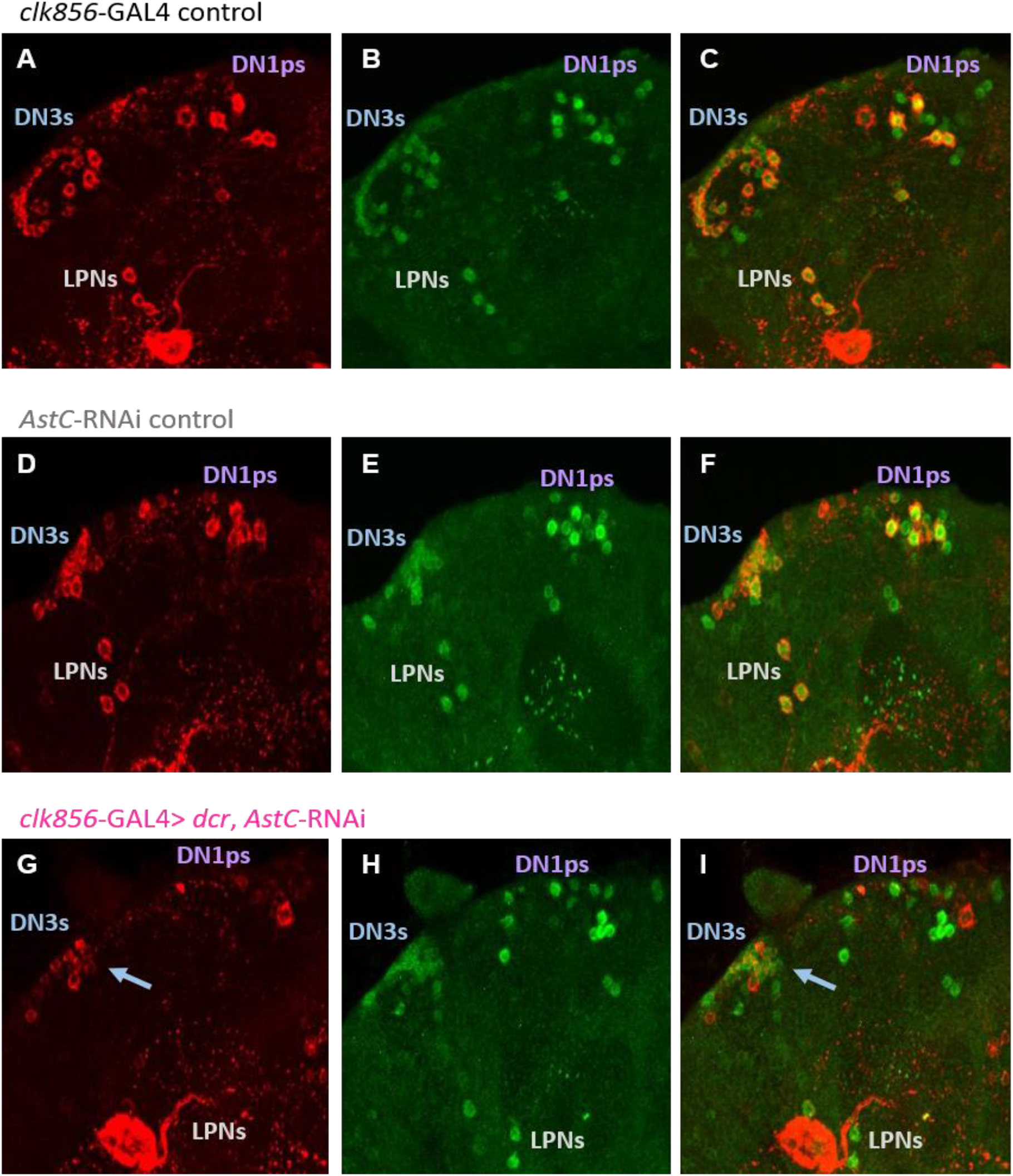
Immunostaining of *CLK856*-GAL4 mediated AstC knock-down reveals ~11-13 DN3s are unaffected. Related to Figure 4 and S5. The GAL4 control (A-C), UAS control (D-F), and the experimental knock-down genotype (G-I) were stained with anti-AstC (red) and anti-TIM (green) at ZT20. ***A***-***F***, For both parental controls, AstC is expressed in four DN1ps, approximately half the DN3s, and the three LPNs. ***G***-***I***, AstC expression is abolished only in the DN1ps and LPNs. Approximately 11-13 DN3s still express AstC, denoted by the arrow. At least eight brains of each genotype were stained together under the same conditions. Confocal settings were maintained the same for all genotypes.

**Figure S7-.**
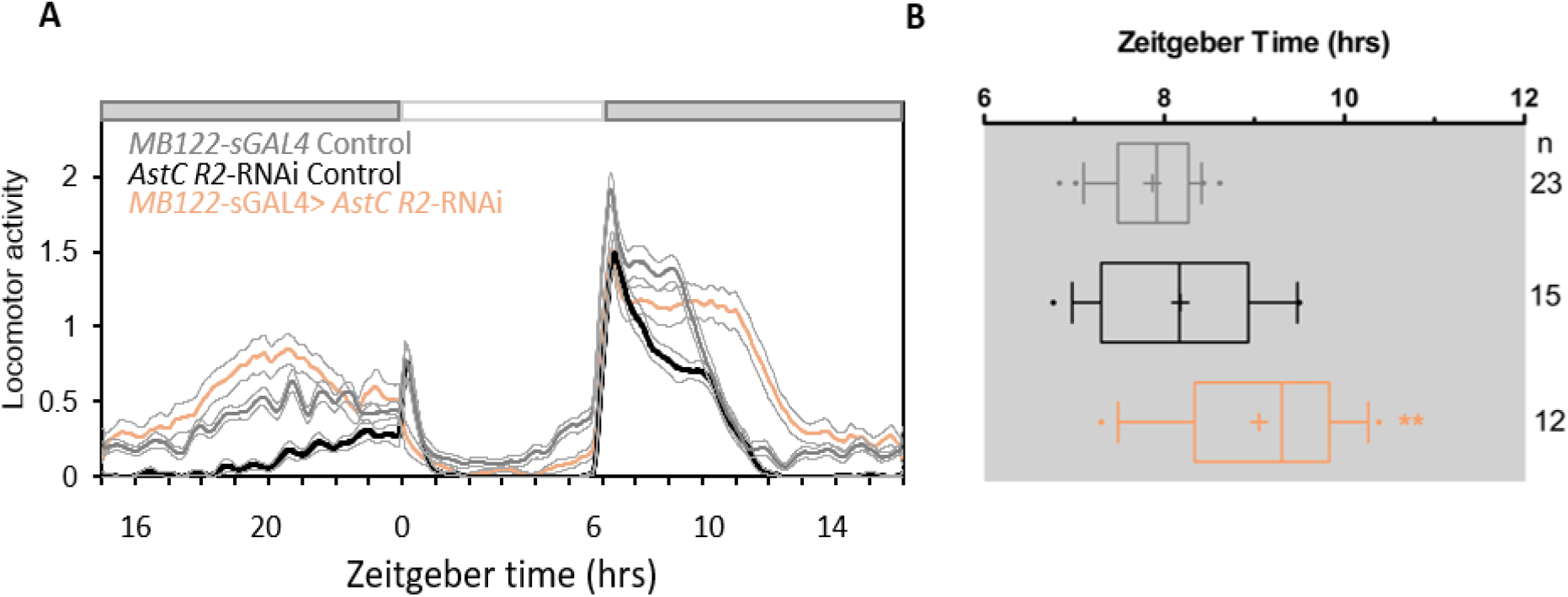
AstC-R2 in LNds is required to regulate evening phase in short photoperiod days. Related to Figure 5. ***A***, Averaged actograms of the *MB122*-sGAL4 control (gray, n=23), *AstC R2-*RNAi control (black, n=15), and the AstC R2-RNAi knock-down in the LNds mediated by *MB122*-sGAL4 (orange, n=12) under short photoperiod 6:18 LD at 27°C. White and dark boxes indicates the respective light and dark phases. Light gray lines represent SEM. ***B***, Boxplot distribution showing the evening peak phase from individual flies. The *MB122*-sGAL4 mediated *AstC R2-*RNAi knock-down (orange) is significantly delayed compared to both the *MB122*-sGAL4 control (gray) and *AstC R2-*RNAi control (black; *p* < 0.001). ** *p* < 0.001. n.s. no significant difference (*p* ≥ 0.05). “+” indicates the mean and the whiskers denotes the 10^th^/90^th^ percentiles.

## References

1. Majercak, J., et al., How a circadian clock adapts to seasonal decreases in temperature and day length. Neuron, 1999. 24(1): p. 219–30.

2. Yoshii, T., D. Rieger, and C. Helfrich-Forster, Two clocks in the brain: an update of the morning and evening oscillator model in Drosophila. Prog Brain Res, 2012. 199: p. 59–82.

3. Grima, B., et al., Morning and evening peaks of activity rely on different clock neurons of the Drosophila brain. Nature, 2004. 431(7010): p. 869–73.

4. Stoleru, D., et al., Coupled oscillators control morning and evening locomotor behaviour of Drosophila. Nature, 2004. 431(7010): p. 862–8.

5. Renn, S.C., et al., A pdf neuropeptide gene mutation and ablation of PDF neurons each cause severe abnormalities of behavioral circadian rhythms in Drosophila. Cell, 1999. 99(7): p. 791–802.

6. Peng, Y., et al., Drosophila Free-Running Rhythms Require Intercellular Communication. PLoS Biol, 2003. 1(1): p. E13.

7. Lin, Y., G.D. Stormo, and P.H. Taghert, The neuropeptide pigment-dispersing factor coordinates pacemaker interactions in the Drosophila circadian system. J Neurosci, 2004. 24(36): p. 7951–7.

8. Hermann, C., et al., Neuropeptide F immunoreactive clock neurons modify evening locomotor activity and free-running period in Drosophila melanogaster. J Comp Neurol, 2012. 520(5): p. 970–87.

9. Johard, H.A., et al., Peptidergic clock neurons in Drosophila: ion transport peptide and short neuropeptide F in subsets of dorsal and ventral lateral neurons. J Comp Neurol, 2009. 516: p. 59–73.

10. Hermann-Luibl, C., et al., The ion transport peptide is a new functional clock neuropeptide in the fruit fly Drosophila melanogaster. J Neurosci, 2014. 34(29): p. 9522–36.

11. Shafer, O.T., et al., Reevaluation of Drosophila melanogaster’s neuronal circadian pacemakers reveals new neuronal classes. J Comp Neurol, 2006. 498(2): p. 180–93.

12. Kunst, M., et al., Calcitonin gene-related peptide neurons mediate sleep-specific circadian output in Drosophila. Curr Biol, 2014. 24(22): p. 2652–64.

13. Goda, T., et al., Drosophila DH31 Neuropeptide and PDF Receptor Regulate Night-Onset Temperature Preference. J Neurosci, 2016. 36(46): p. 11739–11754.

14. Abruzzi, K.C., et al., RNA-seq analysis of Drosophila clock and non-clock neurons reveals neuron-specific cycling and novel candidate neuropeptides. PLoS Genet, 2017. 13(2): p. e1006613.

15. Kreienkamp, H.J., et al., Functional annotation of two orphan G-protein-coupled receptors, Drostar1 and #x2212;2, from Drosophila melanogaster and their ligands by reverse pharmacology. J Biol Chem, 2002. 277(42): p. 39937–43.

16. Price, M.D., et al., Drosophila melanogaster flatline encodes a myotropin orthologue to Manduca sexta allatostatin. Peptides, 2002. 23(4): p. 787–94.

17. Myers, M.P., et al., Light-induced degradation of TIMELESS and entrainment of the Drosophila circadian clock. Science, 1996. 271: p. 1736–1740.

18. Zeng, H., et al., A light-entrainment mechanism for the Drosophila circadian clock. Nature, 1996. 380: p. 129–135.

19. Zitnan, D., F. Sehnal, and P.J. Bryant, Neurons producing specific neuropeptides in the central nervous system of normal and pupariation-delayed Drosophila. Dev Biol, 1993. 156(1): p. 117–35.

20. Guo, F., et al., Circadian neuron feedback controls the Drosophila sleep–activity profile. Nature, 2016. 536(7616): p. 292–7.

21. Rieger, D., R. Stanewsky, and C. Helfrich-Forster, Cryptochrome, compound eyes, Hofbauer-Buchner eyelets, and ocelli play different roles in the entrainment and masking pathway of the locomotor activity rhythm in the fruit fly Drosophila melanogaster. J Biol Rhythms, 2003. 18(5): p. 377–91.

22. Lu, B., et al., Circadian modulation of light-induced locomotion responses in Drosophila melanogaster. Genes Brain Behav, 2008. 7(7): p. 730–9.

23. Gummadova, J.O., G.A. Coutts, and N.R. Glossop, Analysis of the Drosophila Clock promoter reveals heterogeneity in expression between subgroups of central oscillator cells and identifies a novel enhancer region. J Biol Rhythms, 2009. 24(5): p. 353–67.

24. Liang, X., T.E. Holy, and P.H. Taghert, A Series of Suppressive Signals within the Drosophila Circadian Neural Circuit Generates Sequential Daily Outputs. Neuron, 2017. 94(6): p. 1173–1189 e4.

25. Liang, X., T.E. Holy, and P.H. Taghert, Synchronous Drosophila circadian pacemakers display nonsynchronous Ca(2)(+) rhythms in vivo. Science, 2016. 351(6276): p. 976–81.

26. Helfrich-Forster, C., The neuroarchitecture of the circadian clock in the brain of Drosophila melanogaster. Microsc Res Tech, 2003. 62(2): p. 94–102.

27. Birgul, N., et al., Reverse physiology in drosophila: identification of a novel allatostatin-like neuropeptide and its cognate receptor structurally related to the mammalian somatostatin/galanin/opioid receptor family. Embo j, 1999. 18(21): p. 5892–900.

28. Lenz, C., M. Williamson, and C.J. Grimmelikhuijzen, Molecular cloning and genomic organization of a second probable allatostatin receptor from Drosophila melanogaster. Biochem Biophys Res Commun, 2000. 273(2): p. 571–7.

29. Shafer, O.T., et al., Widespread receptivity to neuropeptide PDF throughout the neuronal circadian clock network of Drosophila revealed by real-time cyclic AMP imaging. Neuron, 2008. 58(2): p. 223–37.

30. Williamson, M., et al., Molecular cloning, genomic organization, and expression of a C-type (Manduca sexta-type) allatostatin preprohormone from Drosophila melanogaster. Biochem Biophys Res Commun, 2001. 282(1): p. 124–30.

31. Veenstra, J.A., Allatostatin C and its paralog allatostatin double C: The arthropod somatostatins. Insect Biochemistry and Molecular Biology, 2009. 39(3): p. 161–170.

32. Veenstra, J.A., Allatostatins C, double C and triple C, the result of a local gene triplication in an ancestral arthropod. General and Comparative Endocrinology, 2016. 230-231: p. 153–157.

33. Tanaka, M., et al., Somatostatin neurons form a distinct peptidergic neuronal group in the rat suprachiasmatic nucleus: a double labeling in situ hybridization study. Neurosci Lett, 1996. 215(2): p. 119–22.

34. Fukuhara, C., et al., Phase advances of circadian rhythms in somatostatin depleted rats: effects of cysteamine on rhythms of locomotor activity and electrical discharge of the suprachiasmatic nucleus. J Comp Physiol A, 1994. 175(6): p. 677–85.

35. Biemans, B.A., M.P. Gerkema, and E.A. Van der Zee, Increase in somatostatin immunoreactivity in the suprachiasmatic nucleus of aged Wistar rats. Brain Res, 2002. 958(2): p. 463–7.

36. Funk, C.M., et al., Role of Somatostatin-Positive Cortical Interneurons in the Generation of Sleep Slow Waves. J Neurosci, 2017. 37(38): p. 9132–9148.

37. Dulcis, D., et al., Neurotransmitter switching in the adult brain regulates behavior. Science, 2013. 340(6131): p. 449–53.

38. Deats, S.P., W. Adidharma, and L. Yan, Hypothalamic dopaminergic neurons in an animal model of seasonal affective disorder. Neurosci Lett, 2015. 602: p. 17–21.

39. Dumbell, R.A., et al., Somatostatin Agonist Pasireotide Promotes a Physiological State Resembling Short-Day Acclimation in the Photoperiodic Male Siberian Hamster (Phodopus sungorus). J Neuroendocrinol, 2015. 27(7): p. 588–99.

40. Kaneko, M. and J.C. Hall, Neuroanatomy of cells expressing clock genes in Drosophila: Transgenic manipulation of the period and timeless genes to mark the perikarya of circadian pacemaker neurons and their projections. The Journal of Comparative Neurology, 2000. 422(1): p. 66–94.

41. Zhang, Y., et al., Light and temperature control the contribution of specific DN1 neurons to Drosophila circadian behavior. Curr Biol, 2010. 20(7): p. 600–5.

42. Freeman, M., Reiterative use of the EGF receptor triggers differentiation of all cell types in the Drosophila eye. Cell, 1996. 87(4): p. 651–60.

43. Bahn, J.H., G. Lee, and J.H. Park, Comparative analysis of Pdf-mediated circadian behaviors between Drosophila melanogaster and D. virilis. Genetics, 2009. 181(3): p. 965–75.

44. Guo, F., X. Chen, and M. Rosbash, Temporal calcium profiling of specific circadian neurons in freely moving flies. Proc Natl Acad Sci U S A, 2017. 114(41): p. E8780–E8787.

45. Abruzzi, K., et al., RNA-seq Profiling of Small Numbers of Drosophila Neurons. Methods Enzymol, 2015. 551: p. 369–86.

46. Long, X., et al., Quantitative mRNA imaging throughout the entire Drosophila brain. Nat Methods, 2017. 14(7): p. 703–706.

47. Schlichting, M. and C. Helfrich-Forster, Photic entrainment in Drosophila assessed by locomotor activity recordings. Methods Enzymol, 2015. 552: p. 105–23.

